# Optimized ICA-based removal of ocular EEG artifacts from free viewing experiments

**DOI:** 10.1101/446955

**Authors:** Olaf Dimigen, Humboldt-Universität zu Berlin

## Abstract

Combining EEG with eye-tracking is a promising approach to study neural correlates of natural vision, but the resulting recordings are also heavily contaminated by activity of the eye balls, eye lids, and extraocular muscles. While Independent Component Analysis (ICA) is commonly used to suppress these ocular artifacts, its performance under free viewing conditions has not been systematically evaluated and many published findings display residual artifacts. Here I evaluated and optimized ICA-based correction for two tasks with unconstrained eye movements: visual search in images and sentence reading. In a first step, four parameters of the ICA pipeline were systematically varied: the (1) high-pass and (2) low-pass filter applied to the training data, (3) the proportion of training data containing myogenic saccadic spike potentials (SP), and (4) the threshold for eye tracker-based component rejection. In a second step, the eye-tracker was used to objectively quantify correction quality of each ICA solution, both in terms of undercorrection (residual artifacts) and overcorrection (removal of neurogenic activity). As a benchmark, results were compared to those obtained with an alternative spatial filter, Multiple Source Eye Correction (MSEC). With commonly used settings, Infomax ICA not only left artifacts in the data of both tasks, but also distorted neurogenic activity during eye movement-free intervals. However, correction could be drastically improved by training the ICA on optimally filtered data in which SPs were massively overweighted. With optimized procedures, ICA removed virtually all artifacts, including the SP and its associated spectral broadband artifact, with little distortion of neural activity. It also outperformed MSEC in terms of SP correction. Matlab code is provided.

**Author Note:** I would like to acknowledge Maarten De Schuymer, who conducted early explorations of the effects of high-pass filtering that helped to initiate the current work. I am also grateful to Lisa Spiering for assisting with the MSEC correction and Werner Sommer for providing an excellent working environment. Collection of one of the datasets was supported by a grant from DFG (FG-868-A2). Comments or corrections are highly appreciated.

---

Humans actively explore their environment with 2-4 saccadic eye movements per second, or more than 10,000 during every waking hour. Although natural vision is fundamentally trans-saccadic, procedures in electroencephalographic (EEG) research have traditionally aimed to minimize oculomotor behavior by requiring sustained visual fixation. In recent years, however, there has been rising interest in measuring brain-electric activity also during unconstrained viewing situations such as reading (e.g. Dimigen, Sommer, Hohlfeld, Jacobs, & Kliegl, 2011; Henderson, Luke, Schmidt, & Richards, 2013), scene viewing (e.g. Nikolaev, Nakatani, Plomp, Jurica, & van Leeuwen, 2011; Ossandon, Helo, Montefusco-Siegmund, & Maldonado, 2010; Simola, Torniainen, Moisala, Kivikangas, & Krause, 2013), visual search (e.g. Brouwer, Reuderink, Vincent, van Gerven, & van Erp, 2013; Kamienkowski, Ison, Quiroga, & Sigman, 2012; Körner et al., 2014) or whole-body motion (Soto et al., 2018).

With this approach, eye movements are co-recorded with the EEG and the signal is aligned to the beginning or end of spontaneously occurring eye movements, yielding saccade-or fixation-related potentials (SRPs/FRPs), respectively. Fixations onsets, in particular, provide natural time-locking points to study attentional, cognitive, or affective processes during natural vision, since every fixation triggers a renewed sequence of lambda waves (Evans, 1953; Gaarder, Krauskopf, Graf, Kropfl, & Armington, 1964; Yagi, 1979), primarily visually-evoked potentials that share many features with those elicited by passive retinal stimulation (Dandekar, Ding, Privitera, Carney, & Klein, 2012; Kazai & Yagi, 2003; Kornrumpf, Niefind, Sommer, & Dimigen, 2016; Marton, Szirtes, & Breuer, 1985).

Despite their promises, recordings during natural vision are also complicated by serious data-analytical challenges (Baccino, 2011; Dimigen et al., 2011; Nikolaev, Meghanathan, & van Leeuwen, 2016), the most obvious of which are the massive voltage distortions produced by rotation of the eye balls, movements of the eye lids, and contraction of the extraocular muscles (Berg & Scherg, 1991; Keren, Yuval-Greenberg, & Deouell, 2010; Picton et al., 2000; Plöchl, Ossandón, & König, 2012). These ocular artifacts pose inferential hazards because they are not only magnitudes larger than the event-related neural signals, but typically also correlated to experimental condition (due to condition differences in average saccade size, saccade orientation, and fixation duration). Their complete removal is therefore crucial to avoid misinterpretations.

A multitude of methods has been proposed for ocular correction, including those based on EEG-to-EOG regression, dipole modelling/beamforming, PCA, and other variants of blind source separation (Brunia et al., 1989; Croft & Barry, 2000; Delorme, Sejnowski, & Makeig, 2007; Gratton, 1998; Ille, Berg, & Scherg, 2002). Of these, Independent Component Analysis (ICA) is now perhaps most commonly used to remove occasional saccade and blink artifacts in steady-fixation experiments (Delorme et al., 2007; Jung et al., 2000). ICA employs higher order statistics to decompose the EEG into independent components (ICs), linear mixtures of the scalp channels weighted to be maximally temporally independent (Stone, 2004). For this purpose, ICA is typically trained on a portion of the recording – the *training data* – containing a sample of the neural and non-neural sources active during the task. Only ICs believed to reflect neural sources are then back-projected to the electrode space, yielding an artifact-corrected EEG.

Although ICA is now also frequently applied in SRP/FRP studies, its performance on heavily contaminated free viewing data has not been systematically evaluated, nor compared to that of alternative methods. Yet there are reasons to suspect that the quality of ICA-based ocular correction may be overestimated in practice, in particular for free viewing applications. The first is that the corrected data is rarely analyzed time-locked to the saccade itself, but often inspected in continuous data (where small residual artifacts are virtually impossible to spot) or in fixation onset-aligned averages (in which saccade onset-locked artifacts are jittered due to variance in saccade duration). Second, many free viewing studies aggregate across saccades of different orientations, meaning that the most salient corneoretinal (CR) artifacts cancel out (partially) in the average, making it again difficult to detect remaining artifacts at the single-trial level. Third, the vast majority of SRP/FRP studies so far has focused on posterior sites, where residual artifacts are less obvious. In contrast, facial electrooculogram (EOG) electrodes, which show artifacts most clearly, are often excluded before ICA or not plotted. Finally, few studies have tested whether correction removes legitimate neurogenic activity.

If the corrected EEG is instead aligned to the onsets of saccades of a single orientation and inspected at frontal sensors, results are often sobering. Eye muscle-generated saccadic spike potentials (SPs, see below), in particular, are difficult to model with ICA and many existing reports display clear residual SP artifacts in the plotted waveforms (for arbitrarily picked examples see Dimigen, Kliegl, & Sommer, 2012; Henderson et al., 2013; Kamienkowski et al., 2012; Körner et al., 2014; Kornrumpf et al., 2016; Metzner, von der Malsburg, Vasishth, & Rösler, 2015; Nikolaev et al., 2011; Ossandon et al., 2010; Simola et al., 2013).

The presence of residual artifacts in most published free viewing studies indicates that correction procedures need to be specifically adapted for this purpose. In case of ICA, this requires addressing three practical problems: How to (1) select and preprocess the training data, (2) how to reliably categorize ICs as ocular/non-ocular, and (3) how to measure and minimize the distortion of neural activity. The preprocessing of the input data, in particular, has received comparatively little attention, but has been shown to be more important than the choice of ICA algorithm (e.g. Infomax, FastICA, or AMICA; Zakeri et al., 2014). Rather than proposing yet another correction method, goal of the present study was therefore to validate and adapt the pipeline for the widely-used extended Infomax algorithm (Bell & Sejnowski, 1995; Lee, Girolami, & Sejnowski, 1999) to natural viewing.

In the following, I will first briefly review the three partially independent mechanisms generating ocular artifacts. I will then describe the four manipulated parameters of the ICA pipeline. Finally, I will outline how parallel eye-tracking can be used to more objectively quantify correction outcomes.

## Three types of ocular artifacts

### Corneoretinal (CR) dipole

Due to metabolic activity in the pigmented layer of the retina (Marmor & Zrenner, 1993), each eye ball posesses an electrical gradient of one to several millivolts (depending on ambient illumination level, Marmor & Zrenner, 1993; Young & Sheena, 1988) between its front (cornea) and back. When the eyes rotate, these corneoretinal dipoles also rotate, such that a rightward saccade, for example, generates maximal positive distortions over frontolateral right-hemispheric electrodes (towards which the corneas rotate). For saccades up to 30-40° (Shinomiya, Itsuki, Kubo, & Shiota, 2008), CR artifact amplitude increases as a linear function of saccade size (at ~9.5-16 µV per degree; Keren et al., 2010), meaning that propagation factors are similar for small and large saccades of the same orientation. However, artifact topographies are not strictly mirrosymmetric for up- versus downward saccades (Picton et al., 2000; Plöchl et al., 2012) and have been reported to depend on the participant’s absolute screen viewing position (Ai, Sato, Singh, & Wagatsuma, 2016), which of course changes frequently during free viewing.

### Eye lids

During blinks, the eye lids slide across the positive corneas, allowing current to flow to the forehead (Matsuo, Peters, & Reilly, 1975). A smaller blink-like artifact – sometimes called eye “rider artifact” (Lins, Picton, Berg, & Scherg, 1993) – also occurs towards the end of upwards or oblique-upwards saccades (Barry & Jones, 1965; Plöchl et al., 2012) and is believed to be caused by the eye lids lagging behind the upwards-rotating eye balls, temporarily changing their overlap with the cornea. The result is a blink-like frontal positivity that begins during upward saccades but outlasts saccade offset (by ~100 ms in Plöchl et al., 2012).

### Spike potential

The most difficult-to-correct artifact is the saccadic spike potential (SP), a brief, high-frequency biphasic wave, which ramps up ~5-10 ms before the saccade and reaches its primary peak at saccade onset (Blinn, 1955; Keren et al., 2010; see Carl et al., 2012 for MEG). The main peak of the SP has a focal negative maximum at facial electrodes but is accompanied by more widespread positive deflection over parietal sites. SP topography also changes with saccade direction, but lateralization of its main peak is opposite to that of CR artifacts such that for a rightward saccade, the frontally-negative main spike is largest near the right eye, whereas the diffuse parietal pole shifts towards the left hemisphere. However, the SP is less lateralized than the CR artifact around the eyes, which explains why bipolar EOG montages do not capture this artifact well. Instead, it is largest in a “radial” EOG montage for which facial electrodes are referenced against a centroparietal site. SP amplitude increases with saccade size, although it is unclear whether this relationship is linear (Keren et al., 2010) or not (Boylan & Ross Doig, 1989).

Current scientific consensus holds that the SP most likely reflects myogenic (EMG) activity from the recruitment and only initially synchronous firing of motor units of the extraocular muscles at saccade onset (Thickbroom & Mastaglia, 1985; Yamazaki, 1968). The SP is not of corneoretinal origin since it precedes rotation of the eye ball and survives removal of the bulbus in patients with an eye ball prothesis but preserved eye muscles (Thickbroom & Mastaglia, 1985b). A myogenic rather than cerebral source is also suggested by its frontal generators (Hipp & Siegel, 2013), its presence in darkness (Riggs, Merton, & Morton, 1974) and intramuscular EMG (Yamazaki, 1968) as well as its weakness in intracranial EEG (Jerbi et al., 2009; Kovach et al., 2011; Sakamoto, Lüders, & Burgess, 1991) and patients with extraocular muscle palsy (Thickbroom & Mastaglia, 1986). In contrast, there is little persuasive evidence for cerebral contributions to the SP itself (Balaban & Weinstein, 1985; Berchicci, Stella, Pitzalis, Spinelli, & Di Russo, 2012; Parks & Corballis, 2008), although relevant cortical activity may of course take place during the same interval (e.g. Parks & Corballis, 2008).

Spike potentials have received attention because even involuntary microsaccades (< 1°) during attempted fixation generate sizeable SPs (Armington, 1978; Dimigen, Valsecchi, Sommer, & Kliegl, 2009; Yamazaki, 1968; Yuval-Greenberg, Tomer, Keren, Nelken, & Deouell, 2008), which introduce a broad band artifact in the time-frequency spectrum of the EEG, affecting the low-amplitude beta and gamma bands, in particular (Reva & Aftanas, 2004; Yuval-Greenberg et al., 2008). Complete removal of SPs with ICA has proven challenging even for microsaccades (Craddock, Martinovic, & Müller, 2016; Hassler, Barreto, & Gruber, 2011; Hipp & Siegel, 2013; Keren et al., 2010) and ICA often fails to single out the SP in one or more distinct ICs (Hipp & Siegel, 2013; Keren et al., 2010). In addition, even clean SPs components can be difficult to spot in scalp maps (inverse weights), especially if EOG electrodes were not placed around both eyes, recorded with a bipolar montage, or excluded before ICA.

Residual SP artifacts also pose problems for SRP/FRP analyses in free viewing. First, they distort the typical baseline period in the time and frequency domain. Placing the baseline further away from saccade onset (e.g. +200 to +100 ms) is possible (Nikolaev et al., 2016; Simola et al., 2013), but will reduce the signal-to-noise ratio of the averaged waveforms. The spectral broadband artifact can also leak into neighboring time windows in time-frequency analyses. Second, and more importantly, the SP greatly complicate the study of phenomena occurring in temporal proximity to saccade onset, such as remapping (Kusunoki & Goldberg, 2003), saccadic suppression (Duffy & Lombroso, 1968), or changes in cortical excitability (Ito, Maldonado, Singer, & Grün, 2011). Finally, because a new saccade is executed every 200-400 ms during natural vision, late intervals of the SRP/FRP waveform are contaminated by SPs from the following saccade on the stimulus (see also Figure 6). It is therefore desirable to fully remove SPs.

## Explored parameters of the ICA pipeline

### High-pass filter

The reliability of ICA decompositions has been shown to improve after signal offsets are removed by subtracting the mean voltages across each epoch (mean-centering, Groppe, Makeig, & Kutas, 2009). In addition, practical experience suggests that decompositions improve if slow oscillations and drift are further suppressed by high-pass filtering (Viola, Debener, Thorne, & Schneider, 2010; Winkler, Debener, Muller, & Tangermann, 2015, see also Miyakoshi, 2018; Zakeri, Assecondi, Bagshaw, & Arvanitis, 2014). The adverse effects of slow signals on unmixing quality are not fully understood (Viola et al., 2010; Winkler et al., 2015), but one likely reason is that ICA is biased towards these high-amplitude signals since it tends to focus on data expressing the most power. Filtering may also help to satisfy ICA’s spatial stationarity assumption by removing the slow and spatially unstable changes from skin potential fluctuations and electrode potentials.

Using data from an auditory oddball task, Winkler et al. (2015) systematically investigated effects of high-pass filtering on artifact reduction. Unmixing weights were computed on filtered data and then applied to the unfiltered recording (to preserve slow ERPs like P300, Viola et al., 2010). Filtering at 1 or 2 Hz produced the most dipolar ICs (Delorme, Palmer, Onton, Oostenveld, & Makeig, 2012), the best discrimination between targets/non-targets, and the least noisy ERPs. While these results underline the importance of data preprocessing, adequate filtering may be even more important for free viewing applications. The reason is that during normal vision – and in contrast to tasks with isolated saccades (e.g. Plöchl et al., 2012) – CR artifacts from multiple saccades frequently sum up to produce large deviations from baseline. This problem is most obvious in reading, where ~85% of all saccades point in the same direction (Rayner, 1998), creating DC offsets of ±250 µV at the end of each trial (Dimigen et al., 2011; their Figure 1C). To explore this hypothesis, I high pass-filtered the training data at 20 different low cutoffs.

### Low-pass filter

Removing high frequencies can also improve decompositions since it attenuates electromagnetic noise and scalp-EMG; cutoffs around 40-45 Hz are therfore commonly applied to ICA input data (e.g. Castellanos & Makarov, 2006; Gwin, Gramann, Makeig, & Ferris, 2010; Mannan, Kim, Jeong, & Kamran, 2016; Winkler et al., 2015; Zakeri et al., 2014). However, EMG is not only produced by face, head, and neck muscles, but presumably also reflected in the SP, whose bandwidth extends to at least 90 Hz (Keren et al., 2010; Nativ, Weinstein, & Rosas-Ramos, 1990). This implies that SPs may actually be modelled better if high frequencies remain in the data. To test this hypothesis, the training data was low pass-filtered at 40 or 100 Hz.

### Spike potential overweighting

The difficulty of removing the SP with ICA is likely due to the fact that it accounts for little energy in the signal, because it (1) is of moderate amplitude compared to CR artifacts, (2) possesses a different topography for different saccade directions, (3) lasts only a few samples (~20 ms, see Figure 5), and (4) also reverses its topography within this interval. A potential solution to this problem, suggested by Keren et al. (2010), is to overweight peri-saccadic samples in the training data (see also Mennes, Wouters, Vanrumste, Lagae, & Stiers, 2010). In particular, both Keren et al. (2010) and Craddock et al. (2016) trained their ICAs on relatively short segments (81 ms and 200 ms long, respectively) centered on microsaccades. Similarly, Meyberg et al. (2017) found that only the inclusion of 15° saccades in the training data allowed for the removal of small CR artifacts produced by microsaccades. Finally, to aid component identification, Hassler et al. (2011) proposed an unconventional use of ICA for which “virtual” channels are added to the data, which contain a copy of the SP waveform (averaged across frontal channels) during microsaccade intervals, but zeros elsewhere. Taken together, these studies suggest that overweighting can improve ocular correction, particularly in case of the SP, but this approach has not been tested on free viewing data. I therefore aimed to improve correction by massively overweighing peri-saccadic samples.

### Eye tracker-guided IC classification

Unlike the EOG, eye-tracking provides an accurate gaze position signal that is electrically independent of the EEG and therefore potentially useful to improve correction procedures (Dimigen et al., 2011; Kierkels, Riani, Bergmans, & van Boxtel, 2007; Lourenço, Abbott, & Faisal, 2016; Mannan et al., 2016; Noureddin, Lawrence, & Birch, 2012; Plöchl et al., 2012).

One application is the objective classification of ocular ICs (Dimigen et al., 2011; Plöchl et al., 2012). A simple but elegant criterion for this purpose was proposed by Plöchl and colleagues (2012), who validated it on data from a guided-saccade task. Basis for their classification is the variability of each IC’s activity time course during saccade versus fixations: ICs showing relatively more variance during saccade intervals are likely to reflect ocular artifacts, whereas those showing more variance during fixations are likely neurogenic (because each fixation evokes lambda waves). Thus, if the ratio of both variances (varsaccade/varfixation) exceeds 1, an IC is likely to reflect artifact; conversely, ratios below 1 indicate neural sources. Non-ocular artifacts such as cardiac activity show similar variance during both intervals and are not flagged; they can be detected with other techniques (Campos Viola et al., 2009; Chaumon, Bishop, & Busch, 2015; Mognon, Jovicich, Bruzzone, & Buiatti, 2011; Nolan, Whelan, & Reilly, 2010; Winkler, Haufe, & Tangermann, 2011).

Obviously, the success of this procedure depends on an appropriate threshold. Although a value of 1 seems like a logical choice, Plöchl et al. (2012) proposed using a slightly higher threshold (1.1) to reduce misclassifications. Here I explored different thresholds to identify the lowest threshold that removes ocular ICs while preserving neurogenic activity (see below).

## Quantifying correction with eye-tracking

Correction can distort data in two ways. If ocular ICs are missed, artifacts will remain in the data (*undercorrection*). A question less often discussed is whether ICA removes brain activity observed in the task (overcorrection; e.g. Castellanos & Makarov, 2006; Mennes et al., 2010; Pontifex, Gwizdala, Parks, Billinger, & Brunner, 2016). Overcorrection can happen if the ICA produces mixed neural/non-neural ICs (e.g. Castellanos & Makarov, 2006; McMenamin et al., 2010) or if the experimenter removes neurogenic ICs. For example, inquiries among colleagues suggest that some labs tend to remove all ICs with a frontal topography that is focal or bipolar around the eyes. Presumably, however, at least some of these sources reflect (pre)frontal brain activity.

Under most circumstances, it is impossible to quantify overcorrection since the “true” (artifact-free) data is unavailable for comparison. Here I argue that high-resolution eye-tracking provides a unique opportunity to test for overcorrection, since it allows us to identify short intervals of data *objectively free* of significant oculomotor activity (Dimigen et al., 2011). In particular, every experiment contains at least some intervals during which the eyes were (virtually) motionless and the eye lids remained open. Although such intervals are rare, they can be found immediately after stimulus onsets, since any sufficiently strong stimulation triggers saccadic inhibition, a transient decrease in the rate of saccades (Reingold & Stampe, 2002) and microsaccades (Engbert & Kliegl, 2003). Since these intervals should not be modified by ocular correction, they provide a ground truth to quantify the overcorrection produced by different methods^1^.

## A benchmark for ICA: Multiple-Source Eye Correction

To put the performance of Infomax ICA into context, I compared it to that obtained with an alternative algorithm, the surrogate variant of *Multiple-Source Eye Correction* (MSEC, Berg & Scherg, 1994). Like ICA, MSEC can be described as a spatial filter (Ille et al., 2002), which separates brain and artifact activities based on their topographical definitions (see *Appendix A*). A main difference to ICA is that artifact topographies are not obtained by blind source separation, but empirically defined by averaging the artifacts of isolated calibration saccades. In addition, a set of generic brain topographies is defined by a dipole model that is the same for all participants. The purpose of this “surrogate” source model is *not* to directly model neural activity in the task, but to reduce the subtraction of brain activity spatially correlated to the artifact topographies (i.e. to reduce overcorrection). Because MSEC has been shown to produce good corneoretinal correction in natural reading (e.g. Dimigen et al., 2011; Kornrumpf et al., 2016) is was considered a suitable benchmark.

## Current study

To summarize, this study aimed to improve and evaluate ICA-based ocular correction of free viewing data. For this purpose, I analyzed combined EEG/eye movement recordings from two frequently studied paradigms with very different oculomotor behavior: visual search in scenes and left-to-right sentence reading. The four described parameters of the ICA pipeline were orthogonally manipulated (see Figure 1). The outcomes of each ICA variant were then compared with objective measures of under- and overcorrection and benchmarked against those obtained with MSEC.

**Figure 1.**
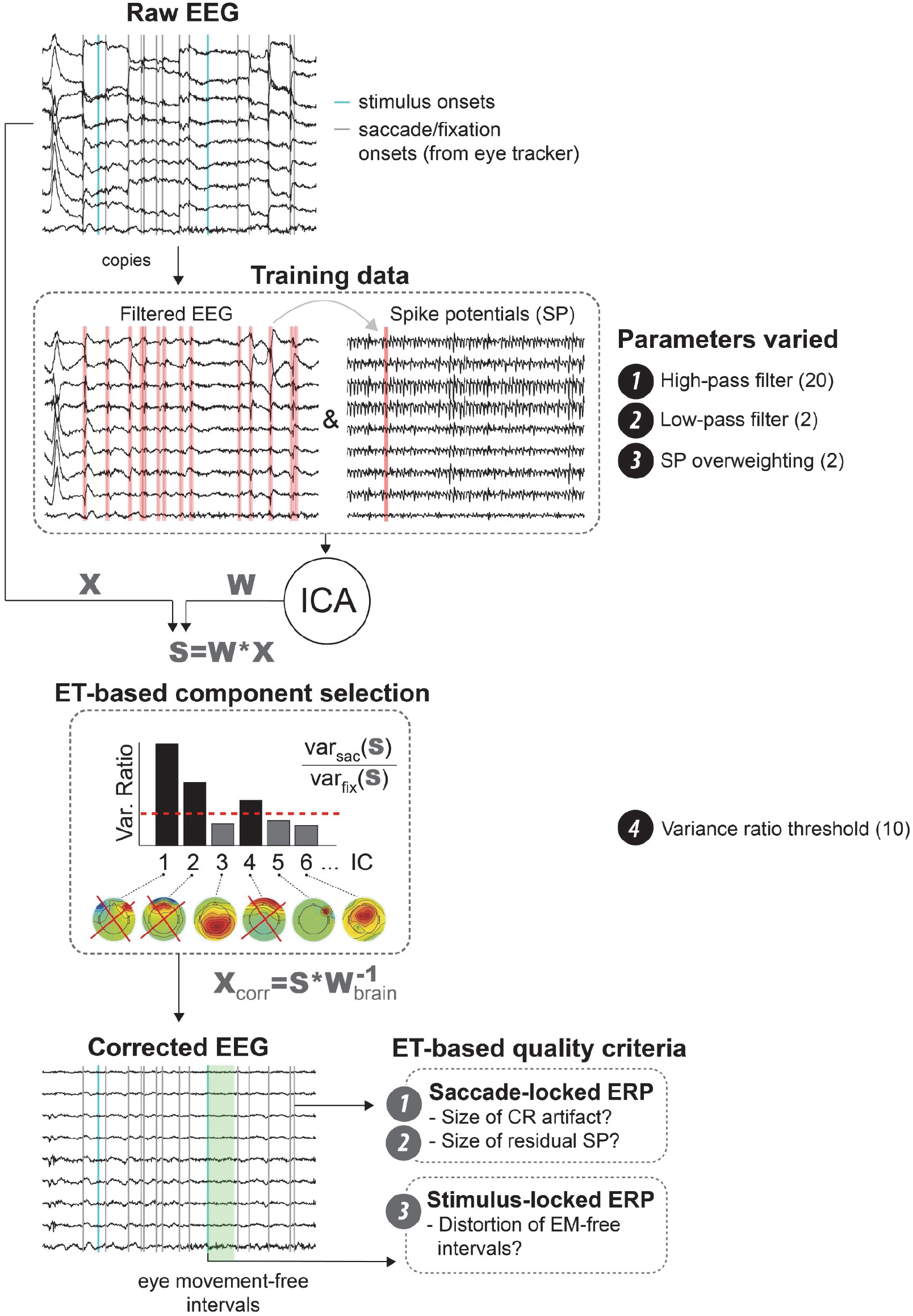
Schematic workflow to determine parameters for optimized ICA training (OPTICAT) and component classification. Steps are described in the section “*Overview over pipeline*”.

## Methods

### Overview over pipeline

I analyzed data from two EEG/eye-tracking experiments with unconstrained eye movements (Figure 2). Both were recorded in the same laboratory with identical hardware and largely identical recording settings (see details below).

Figure 1 summarizes the analysis pipeline. In a first step, I created for each participant 40 differently filtered copies of the original recording by crossing 20 high-pass filters (from 0.016 to 30 Hz) with two low-pass filters (40 or 100 Hz). These differently filtered datasets were then cut into epochs serving as ICA training data. From each of these “basic” versions of the training data, I then created a second, “overweighted” version in which peri-saccadic samples were overrepresented. This was achieved by cutting 30 ms segments around saccade onsets and repeatedly re-appending them to the training data.

In a second step, ICAs were computed on each of the resulting 80 training datasets (20 low cutoffs × 2 high cutoffs × 2 versions). In a third step, the resulting unmixing matrices (**W**) were multiplied with the original, unfiltered recording (**X**), thereby producing activity waveforms (**S**) for the ICs in the original recording. In a fourth step, eye movement events provided by the eye-tracker were used to remove ICs whose activity waveforms showed more variance during saccades than fixations. To find the best threshold for this classification, 10 different thresholds were applied (0.6, 0.7,…1.5), the corresponding ICs removed, and the data back-projected. Since every threshold was applied to every ICA solution, this yielded 800 versions of artifact-corrected data per participant (or 19,200 in total).

Finally, three eye tracker-based criteria were used to compare correction quality. To quantify *undercorrection*, I measured the residual amplitude of (1) CR and (2) SP artifacts in saccade onset-aligned ERPs (SRPs). To quantify *overcorrection*, I identified short stimulus-onset aligned epochs without detectable oculomotor activity. Because these intervals should not be affected by ocular correction, overcorrection was quantified as (3) the degree to which these intervals were changed by ICA.

### Participants

For the current analysis, I used the first 12 participants of each experiment. Most were students at Humboldt University, with a mean age of 25.3 years in the *Scene* (range 19–25 yrs., 7 female) and 21.3 years in the *Reading* experiment (18–33 yrs., 11 female). Procedures complied with the declaration of Helsinki and participants provided written informed consent.

### Scene viewing experiment

In the *Scenes* experiment, participants searched for a target stimulus hidden within greyscale natural images. Most images were taken from the *Zurich Natural Image Database* (Einhäuser, Kruse, Hoffmann, & König, 2006), a set of photographs shot in a forest. During the experiment, 99 images were presented at a resolution of 800 × 600 pixels (28.8° × 21.6°), centered on a 1024 × 768 pixels background showing 1/f noise. On each trial, a single scene was presented and the participant’s task was to find a grey disc (0.4 cd/m^2^) that appeared at a random location within the image 8 to 16 s after scene onset. This disc had an initial diameter of just 0.07° but then slowly increased in size. Once the participant found the target, he/she pressed a button, terminating the trial. The top row of Figure 2 summarizes eye movement behavior in the task. Saccades had a median amplitude of 4.9°, mean duration of 44.2 ms and pointed in all directions, although most were horizontal. Fixation locations were distributed across the images, with some bias towards the image center and corners (heat map in Figure 2). Average fixation duration was 264 ms.

**Figure 2.**
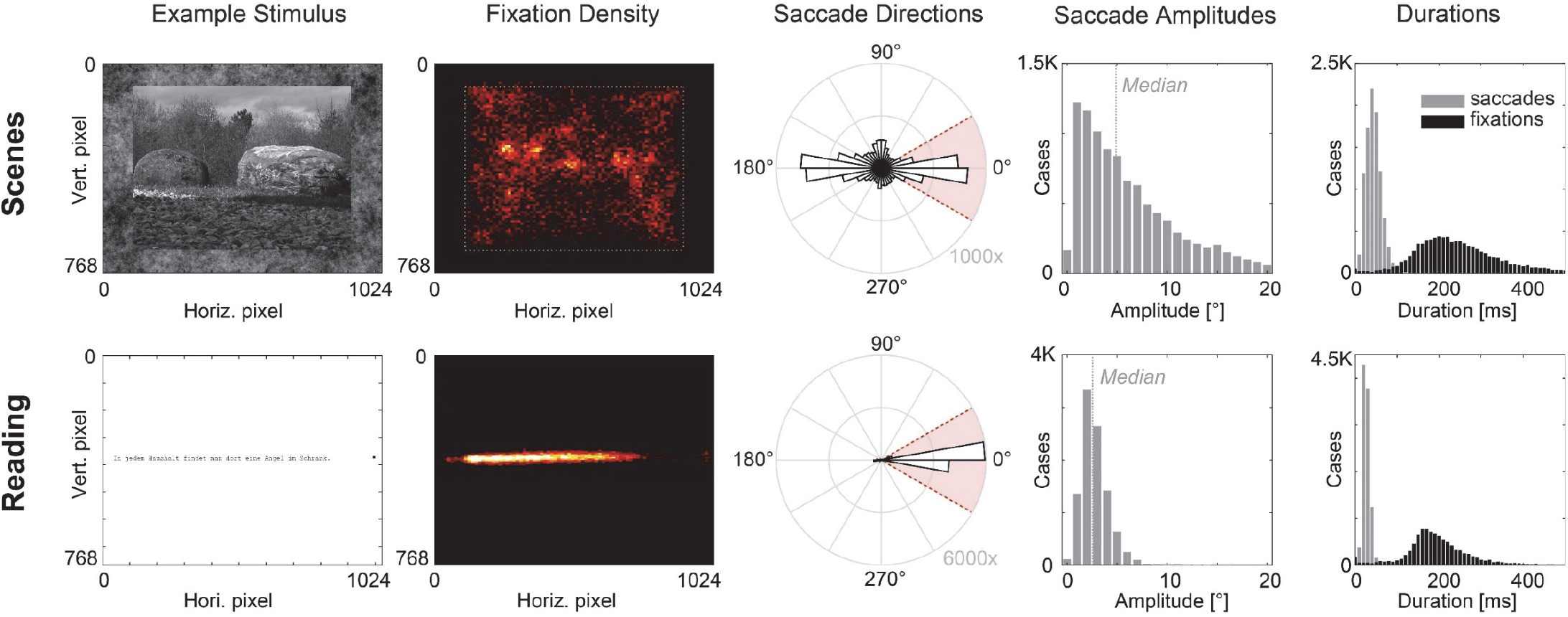
Stimuli and eye movement behavior in the two experiments. In the *Scenes* experiment, participants searched for a small target stimulus hidden within natural images. In the *Reading* experiment, participants read short stories. For each experiment, plots show an example stimulus, a “heat map” of fixation locations (warmer colors indicate higher density), a directional histogram of saccade directions, and the distribution of saccade amplitudes. Red shading in the saccade direction plots indicates saccades classified as “rightward” (±30°).

### Reading experiment

In the *Reading* experiment, participants read 152 short stories (including 8 practice trials) of the Potsdam Sentence Corpus III, a set of sentences used in previous ERP research (Dambacher et al., 2012). Each story consisted of two sentences that were successively presented as single lines of black text (Figure 2) on a white background (0.45° per character). Purpose of the experiment was to study fixation-related N400 effects elicited by a semantically congruent/incongruent word in the second sentence. Only data for the second sentence was analyzed. As Figure 2 shows, reading saccades had a median amplitude of 2.6° and mostly pointed rightward. Average fixation duration was 197 ms. Please note that the relatively short average fixation durations in both experiments (cf. Rayner, 1998) are explained by the sensitive saccade detection algorithm used here, which also detects microsaccades.

### Common methods: Stimulation & Eye-Tracking

Recordings were made in an electromagnetically shielded room. Stimuli were presented on a 22 inch monitor (Iiyama Vision Master Pro 510) at a viewing distance of 60 cm using *Presentation* software (Neurobehavioral Systems Inc.). Binocular eye movements were recorded at a rate of 500 Hz with a video-based IView-X Hi-Speed eye-tracker (SMI GmbH). Offline, saccades and fixations were detected using Engbert & Kliegl’s (2003) velocity-based algorithm as implemented in the EYE-EEG toolbox (Dimigen et al., 2011). Saccades were defined as intervals in which the velocity of both eyes exceeded for ≥ 10 ms a threshold set at 5 median-based SDs of all recorded eye velocities (excluding blink intervals). This low threshold (cf. Dimigen et al., 2009) was chosen to maximize detection sensitivity for microsaccades. In cases where multiple saccades were detected within < 50 ms, I only kept the first to avoid detecting post-saccadic oscillations (Nyström, Hooge, & Holmqvist, 2013) as separate saccades.

### Common methods: Electrophysiology

EEG and EOG were recorded from 45 (*Scenes*) or 63 (*Reading*) Ag/AgCl electrodes referenced against the left mastoid. Electrodes were mounted in a cap at standard 10-10 system positions, except for four EOG electrodes that were affixed to the outer canthus and infraorbital ridge of each eye. Throughout all analyses, EOG electrodes remained in the data and were treated like EEG channels (which is crucial to detect residual artifacts and ocular ICs; Dimigen et al., 2011; Nikolaev et al., 2016). To make the montages in both experiments comparable, I reduced the *Reading* dataset also to 45 electrodes. Signals were recorded with BrainAmp DC amplifiers (Brain Products GmbH) at a rate of 500 Hz with impedances kept < 5 kΩ. The *Scenes* data was initially acquired with a time constant of 10 s, whereas the *Reading* data was acquired as direct current data. To make the datasets directly comparable, the raw data of both experiments was high-pass filtered at 0.016 Hz (10 s time constant) using ERPLAB’s Butterworth filter (*pop_basicfilter.m*, Lopez-Calderon & Luck, 2014). EEG and eye-track were synchronized offline based on shared trigger pulses with a synchronization error < 2 ms.

### Creating differently filtered ICA training data

For each participant, I trained the ICA on 80 differently preprocessed versions of the original data. In a first step, I created 20 copies of the continuous EEG recording and filtered them at different low cutoffs using EEGLAB’s (Delorme & Makeig, 2004) default FIR filter (*pop_eegfiltnew.m* by Andreas Widmann) with its default frequency-dependent transition bandwidth/slope settings (e.g. at 1 Hz: −6 dB attenuation at 0.5 Hz). The low cutoffs used were: 0.016 Hz (no additional filtering), 0.1, 0.25, 0.5, 0.75, 1, 1.5, 2, 2.5, 3, 3.5, 4, 5, 7.5, 10, 12.5, 15, 20, 25, and 30 Hz. Afterwards, each dataset was also low-pass filtered at either 40 or 100 Hz.

Each filtered dataset was then cut into 3 s epochs (−200 to +2800 ms) around stimulus presentations. In the *Scene* dataset, I used the onset of the photograph as well as the onset of the search target, yielding 198 epochs. In the *Reading* data, I used the onset of the second sentence, yielding 152 epochs. Note that these epochs contain a representative sample of brain and artifact activity in these tasks, since they include both the stimulus onset and several subsequent saccades. From each epoch, I then removed the mean channel voltages across the epoch (mean-centering, Groppe et al., 2009). Any high-pass filtering was therefore done in addition to mean-centering. To exclude epochs with large non-ocular artifacts, I rejected epochs containing extreme outliers (> ±500 µV) in any channel. No additional pruning was performed (see also Winkler et al., 2015).

The resulting 40 differently filtered versions of training data will be called “basic” versions in the following, because they contain no overweighted artifacts. From each of these filtered and epoched datasets, I then used the first 162,000 points to train the ICA (corresponding to *k* = 80 points per weight in the 45 × 45 unmixing matrix).

### Overweighting spike potentials

From each basic versions of the training data, I then created an “overweighted” copy in which SP samples were overrepresented (Figure 1). For this purpose, short 30 ms epochs (−20 to +10 ms) were extracted around all saccade onsets found in the basic version. These brief saccade-locked epochs were then again mean-centered, concatenated together, and repeatedly appended to the end of the training dataset until its total length was doubled to 324,000 points (*k* = 160), half of which now only consisted of parts of the SP waveform (i.e. overweighting proportion of 50%). This means that all of the appended samples were already contained in the basic version of the training data, that is, they were redundant except for the renewed mean-centering applied to the brief peri-saccadic epochs.

To test how much overweighting is necessary, I also ran an exploratory analysis in which I varied the amount of overweighting – the number of appended peri-saccadic samples – so that they constituted between 0 and 50% (in steps of 5%) of the overall training data.

### ICA decomposition

ICAs were computed on each of the resulting 80 training datasets using EEGLAB’s binary implementation of extended Infomax ICA (minimum change criterion: 10^-7^). Infomax was chosen because it is widely used as the default option in EEGLAB and produces rather reliable decompositions (Groppe et al., 2009; Pontifex et al., 2016). The resulting ICA weights were then applied to the original, unfiltered recording. More precisely, source waveforms (**S**) for the original recording were computed by multiplying each of the 80 unmixing matrices (**W**) computed on the training data (and the corresponding sphering matrices, stored separately in EEGLAB) with the matrix (**X**) containing the unfiltered recording (Figure 1).

### Eye tracker-guided component identification

The next step was to remove ocular ICs using the procedure by Plöchl et al. as implemented in EYE-EEG (function *pop_eyetrackerica.m*). I applied 10 thresholds between 0.6 and 1.5 (in steps of 0.1). To compute variance ratios, the saccade time window was defined as lasting from −10 ms before saccade onset until saccade offset. Conversely, fixations were defined as lasting from fixation onset until 10 ms before the saccade^2^. To obtain the corrected EEG (**X**_corr_), source waveforms (**S**) from the original recording were multiplied with mixing matrices (**W^-1^_brain_**) in which the ocular ICs were set to zero.

### Measuring correction quality

To quantify correction quality and visualize results, the corrected EEG was converted to average reference. I then extracted three measures of correction quality:

#### Undercorrection: CR artifact size

The size of residual CR artifacts was quantified by averaging all rightward saccades (tolerance of ±30°; see red shading in Figure 2) and measuring the remaining EEG lateralization at frontal electrodes. Rightward saccades were selected because they occur frequently in both scene viewing and reading. However, for *Scenes*, comparable results were obtained for other saccade directions (*Appendix C*). Epochs were cut from −200 to +600 ms around saccade onsets, baseline-corrected from −50 to −10 ms (thereby excluding the SP from the baseline), and averaged. Residual artifact amplitude in these saccade-locked ERPs was then summarized as a single measure by subtracting the mean voltage at eight frontal left-hemispheric EEG and EOG electrodes (LO1, IO1, FP1, AF7, F7, F3, FT9, FC5) from that at their right-hemispheric counterparts (LO2,…, FC6) in the window +10 to +200 ms after saccade onset. Hemispheric lateralization within this window captures the CR artifact but excludes the SP, which was quantified separately.

A possible concern with this measure is that saccades are accompanied by genuine lateralized brain activity. However, lateralized potentials related to attention shifts (Eimer, 2014) or oculomotor preparation (Becker, Hoehne, Iwase, & Kornhuber, 1972; Berchicci et al., 2012; Everling, Krappmann, & Flohr, 1996; Moster & Goldberg, 1990) typically occur *before* rather than after the saccade and most of them are strongest at posterior sites. Brain activity was therefore unlikely to influence post-saccadic frontal lateralization in a relevant manner.

#### Undercorrection: SP size

The same rightward SRPs were used to measure residual SP amplitude. The SP is difficult to capture in a single measure, because it is biphasic and has a diffuse parietal pole. As a suitable measure, I computed the average Global Field Power (GFP, Lehmann & Skrandies, 1980) at the eight sampling points between −8 ms and +8 ms around saccade onset.

#### Overcorrection: Brain signal distortion

Overcorrection was quantified in intervals without significant oculomotor activity. As explained above, a suitable interval to find such intervals is immediately after stimulus onset, i.e. after presentation of the scene/sentence stimulus. Stimulus onset also generates a sequence of visually-evoked potentials (P1 and N1) that should not be altered by ICA in epochs without eye movement. Another set of epochs was therefore cut around stimulus onsets and baseline-corrected with a 100 ms pre-stimulus baseline. The eye-tracker was then used to identify a subset of these epochs in which no blink or eye movement was detected between −100 and +200 ms after stimulus onset; this was the longest feasible interval that yielded some (virtually) eye movement-free epochs for each participant. Three criteria were used to identify these epochs: First, I excluded epochs in which a (micro)saccade was detected. Second, I removed epochs in which gaze position in any of the four eye-tracker channels (vertical and horizontal, left and right eye) changed by > 0.2° between successive samples. Finally, to detect significant binocular drift, I removed epochs during which horizontal or vertical gaze position (averaged binocularly) varied by > 0.5° (peak-to-peak) within the epoch.

These criteria identified *M* = 38.0 eye movement-free epochs per participant for *Scenes* (range: 5-66, SD = 21.1) and *M* = 101.3 (range: 32-129, SD = 26.3) for *Reading* which were then averaged to obtain a “clean” stimulus-ERP. To quantify overcorrection, this ERP was subtracted from the same ERP after ocular correction with each of the 800 ICA variants described above. The resulting difference waves (Figure 7) therefore capture the distortion introduced by ICA and the average GFP of these difference waves across the 300 ms was used to quantify overcorrection.

### Comparison to MSEC

Results were compared to those obtained with surrogate MSEC, as implemented in BESA (version 6.0, Besa GmbH, Gräfeling). Details on the algorithm and its implementation are provided in *Appendix A*.

### Statistics

The three measures of correction quality were entered into separate mixed ANOVAs on the factors *Experiment* (2-level between-subject factor), *High-pass filter* (20), *Low-pass filter* (2), and *Overweighting* (2). For these analyses, the threshold was fixed at 1.1. ANOVAs were conducted using the “ez” package in *R* (Lawrence, 2013). To handle violations of the sphericity assumption, degrees of freedom were adjusted by multiplication with the Greenhouser-Geisser epsilon. Here I report the original degrees of freedom, the epsilon (ε), the adjusted *p*-values, and effect size as generalized eta-squared (ηG^2^). Post-hoc paired *t*-test were used to assess the relationship between threshold and overcorrection and to compare MSEC with the best ICA solution.

## Results

Below, I first present the results for CR and SP correction and for overcorrection. These are based on the removal of ICs with the threshold fixed at 1.1, which proved to be suitable. Afterwards, I summarize the effects of different thresholds. Finally, I compare ICA to MSEC.

### Correction of CR artifacts

Figure 3A summarizes the impact of high-pass filtering, low-pass filtering, and overweighting on the removal of CR artifacts. As evident from this figure, correction quality in both experiments differed drastically as a function of filtering. Results for CR artifacts are also depicted in Figure 4, which shows the grand-average SRP locked to rightward saccades. Panel A in Figure 4 displays the raw SRP without ocular correction; here, strong CR artifacts of up to ±70 µV are evident as differences between left- (blue lines) and right-hemisphere (red lines) channels. Black lines mark sagittal midline electrodes that are less affected by lateral saccades. At these channels, the lambda responses (P1 equivalent) can be seen peaking over occipital cortex about 110 ms after saccade onset.

**Figure 3.**
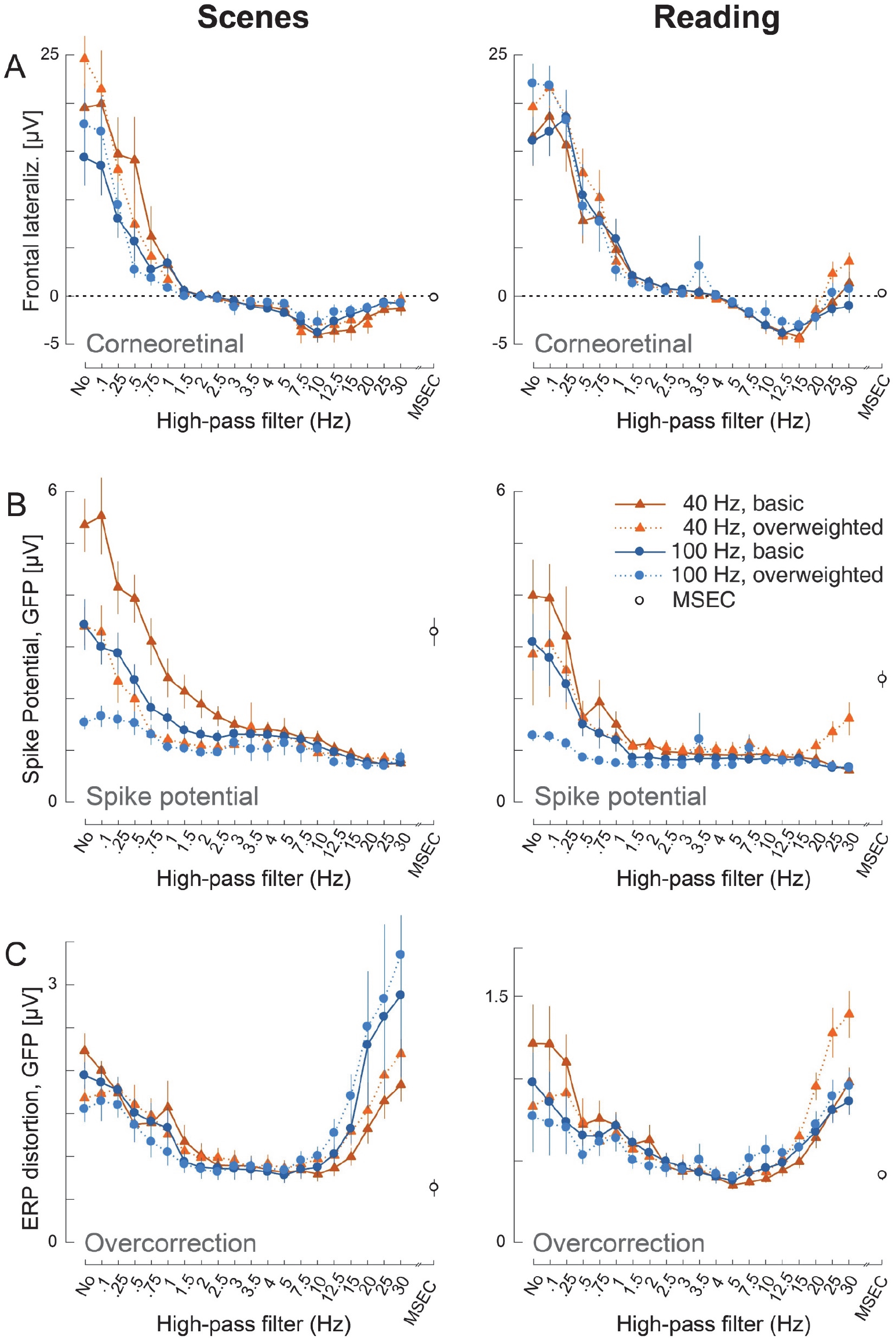
Overview over main results for the *Scenes* (left) and *Reading* (right) experiment. Plots in the three rows depict the (A) size of residual corneoretinal artifact, (B) size of the spike potential and (C) distortion of eye-movement free intervals (overcorrection). These dependent variables are plotted as a function of the *high-pass filter, low-pass filter*, and *overweighting* applied to the ICA training data. For this plot, ocular ICs were flagged using a threshold of 1.1. Results for the MSEC algorithm are shown on the right of each plot (black open circles). Error bars indicate ±1 SEM.

**Figure 4.**
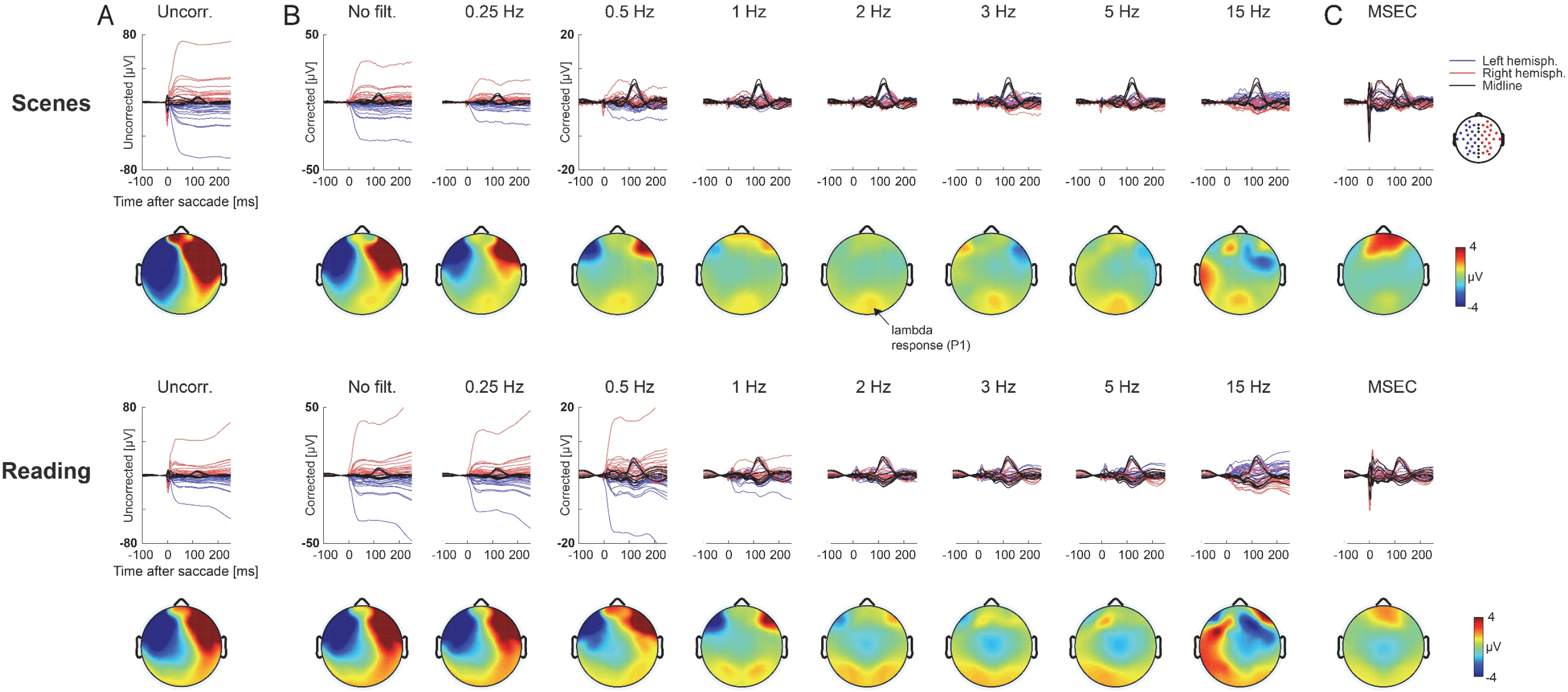
Effect of high-pass filtering the training data on ICA correction of the original, unfiltered data. Plotted is the grand-average saccade-locked ERP (SRP) for rightward saccades in the *Scenes* (upper half) and *Reading* (lower half) experiment. Time zero marks saccade onset. (A) The leftmost column shows the SRP without ocular correction. Strong lateralized CR artifacts are evident. Maps depict the corresponding scalp distribution in the interval 10-200 ms after saccade onset. The positive deflection over midoccipital sites reflects the lambda response. Note that all plots also include EOG electrodes, which show the strongest residual artifacts. (B) Same data corrected with ICA. Optimal results were obtained with high-pass filters of 2-2.5 Hz. Note the different scaling of the y-axis between plots. (C) Same data, corrected with MSEC. (*Note*: results in this figure are based on training data low-pass filtered at 100 Hz, with overweighted SPs)

Figure 4B show the same data after correction with differently trained ICAs. Note that all averages shown here were extracted from the original, *unfiltered* data corrected with ICA; plots only differ in terms of preprocessing of the training data. Although ICA reduced artifacts at all filter settings, visual inspection suggests that residual artifact size depended strongly on the high-pass filter, with cutoffs < 1.5 Hz producing suboptimal correction. Numerically, best results were obtained at cutoffs between 2 and 2.5 Hz for Scenes, and 3-4 Hz for Reading. Interestingly, very high cutoffs (> 5 Hz) tended to invert the topography of residual CR artifacts in the corrected data, that is, right-hemisphere channels became more *negative* than left-hemisphere channels (see values below zero in Figure 3A; see reversed lateralization in Figure 4B). In other words, overly aggressive filtering overcompensated CR artifacts, whereas filtering at < 1.5 Hz left artifacts in the data. This visual impression was confirmed by a significant effect of *high-pass filter* on CR artifact amplitude, *F*(19,418) = 82.917, ɛ = 0.155, *p* < 0.0001, ηG^2^ = 0.638, which did not interact with *Experiment*. Factors *low-pass filter* and *overweighting* did not affect CR correction.

### Correction of spike potential

Figures 3B summarizes results for the SP. Waveforms are depicted in Figure 5. High-pass filtering also strongly influenced SP correction. With an increasing cutoff frequency, residual SP amplitude decreased in an almost monotonous fashion (main effect of *high-pass filter* on residual SP amplitude, *F*(19,418) = 50.940, ɛ = 0.103, *p* < 0.0001, ηG^2^ = 0.380). Importantly, however, the other two parameters also strongly modulated the SP: First, correction was improved by overweighting, but as Figure 3B shows, this benefit was only present in datasets that were only moderately high-pass filtered (at frequencies of about 3 Hz or less), resulting in a significant *high-pass filter* × *overweighing* interaction, *F*(19,418) = 10.943, ɛ = 0.124, *p* < 0.0001, ηG^2^ = 0.086. Second, a similar pattern was seen for the *low-pass filter* (40 vs. 100 Hz). Leaving the filter open up to 100 Hz improved SP correction, but again only if high-pass filtering was moderate (interaction *high-pass filter* × *low-pass filter*, *F*(19,418) = 9.799, ɛ = 0.111, *p* < 0.001, ηG^2^ = 0.068).

**Figure 5.**
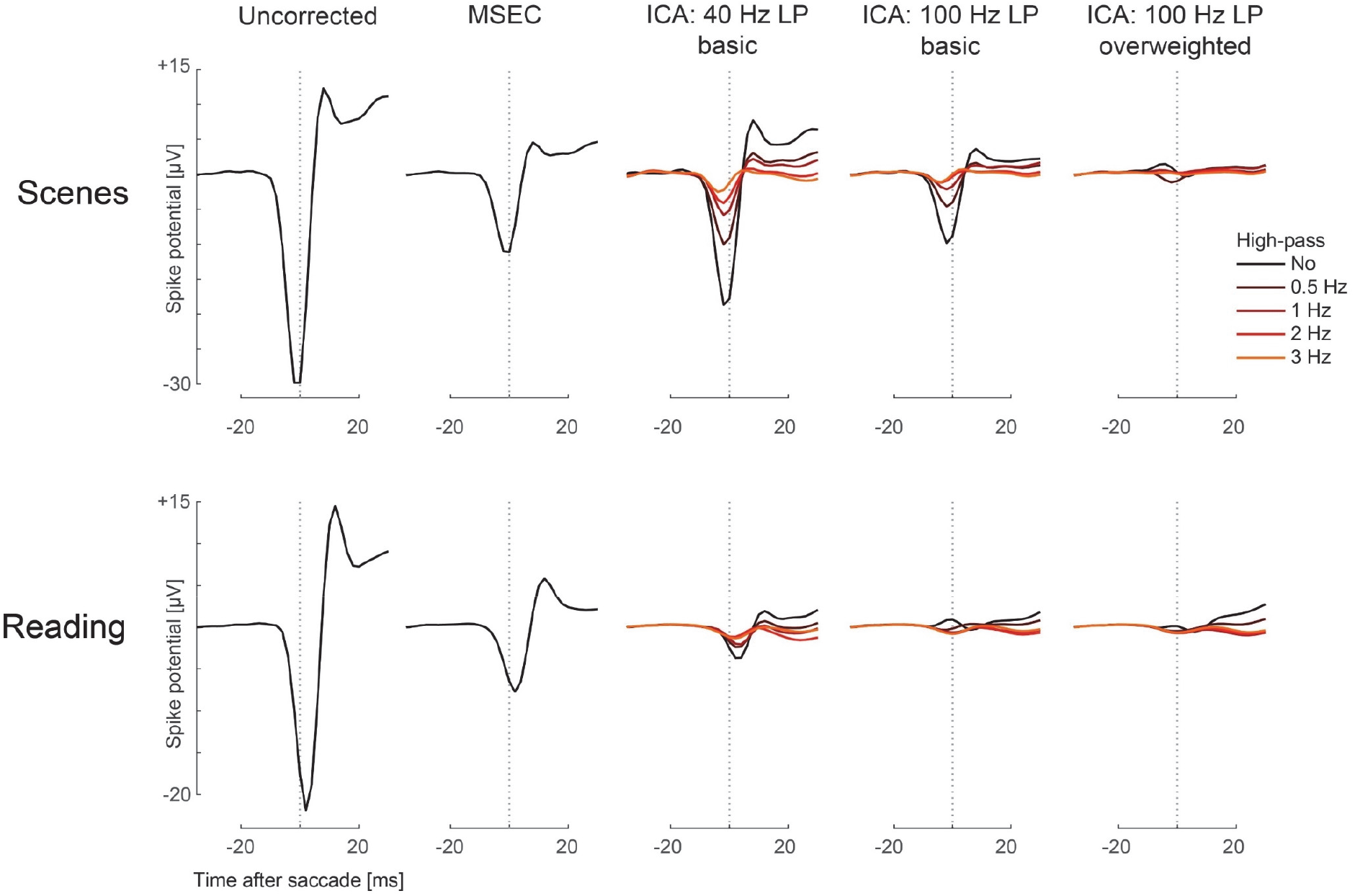
Spike potential correction as a function of *high-pass filter, low-pass filter*, and overweighting. Waveforms show the SRP for rightward saccades at a “radial” EOG channel (mean of all facial EOG electrodes minus electrode POz). Note that the SP is almost entirely removed if the three parameters are combined in an optimal way (rightmost panels). In contrast, MSEC failed to fully correct the SP.

Figure 5 shows that if all three parameters were combined post-hoc in a near-optimal manner, that is, high-pass filtering at 2 Hz, overweighting of SPs, and no additional low-pass filtering, SP artifacts could be virtually eliminated from both *Scenes* and *Reading* data. Figure 6 also shows the Scenes data in the frequency domain, that is, after saccade-locked epochs (cut from −600 to 1000 ms for this analysis) were individually wavelet-transformed using EEGLABs newtimef.m function, yielding event-related spectral pertubation (ERSP) values relative to a pre-saccadic (−200 to −100 ms) spectral baseline. Wheras a more “typically” trained ICA (bandwidth 1-40 Hz, no overweighting) left considerable spectral artifacts in the data, the near-optimally trained ICA removed the broadband artifact in the beta (~14 to 30 Hz) and gamma (> 30 Hz) band (Craddock et al., 2016; Hassler et al., 2011; Hipp & Siegel, 2013; Yuval-Greenberg et al., 2008); not only from the current saccade (at time zero, see Figure 6), but also from the following saccade on the stimulus.

**Figure 6.**
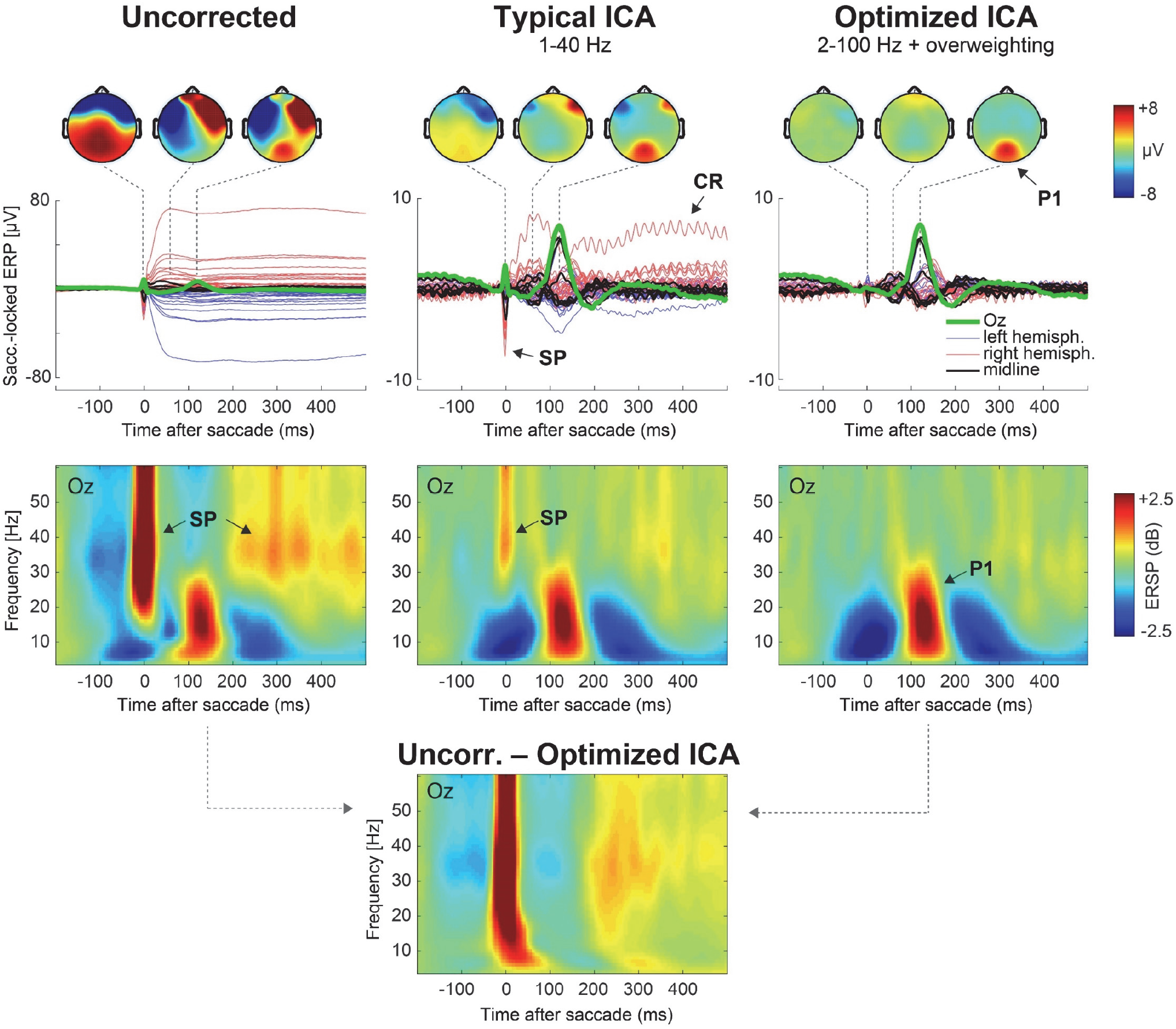
Correction with a “typical” versus “optimized” ICA. In all panels, time zero indicates the onset of rightward saccades during scene viewing. *Left panels*: Grand-average saccade-related activity without correction. *Middle panels*: Same data, corrected with a typically trained ICA (1-40 Hz bandwidth, no overweighting). *Right panels*: Same data, corrected with a well-trained ICA (2-100 Hz, with overweighting). Upper panels plot the grand-average saccade-locked ERPs with occipital electrode Oz highlighted in green. Maps show the scalp distribution at −2 ms (SP artifact), 60 ms (CR artifact) and 120 ms (P1 brain potential), respectively. Lower panels present the same data after a wavelet transform (ERSP). Note how the “typically”-trained ICA leaves artifacts from both the current and next saccade (around 250-300 ms) in the spectrograms, whereas the optimized ICA removed these spectral artifacts. The bottom panel shows the pure artifact in the frequency domain, i.e., the difference between the spectrogram without correction minus those corrected with an optimized ICA. Results for *Reading* are not shown here but were similar.

### Brain signal distortion (overcorrection)

Next, I tested whether ICA distorts neurogenic activity during stimulus-locked intervals (virtually) free of oculomotor activity. Aggregated results are again shown in Figure 3C. Exemplary waveforms are provided in Figure 7 (the data shown is based on ICAs trained on data low-pass filtered at 100 Hz with overweighted SPs, since these settings provided the best SP correction, see above). Panels B and F in Figure 7 show the binocular gaze position in these “clean” epochs, measured by the eye-tracker. In both experiments, grand-average gaze position changed by less than 0.05° across the 300 ms intervals. Panels A and D in Figure 7 depict the corresponding stimulus-ERPs for these “clean” epochs without correction. As expected, the trial-initial stimulation elicited a P1/N1 complex in both experiments. The following panels B and E only show the *difference waves* (at all channels) between this artifact-free ERP *after* ICA correction minus the same ERP *without* any ocular correction. In other words, these panels show the distortions introduced by ICA.

**Figure 7.**
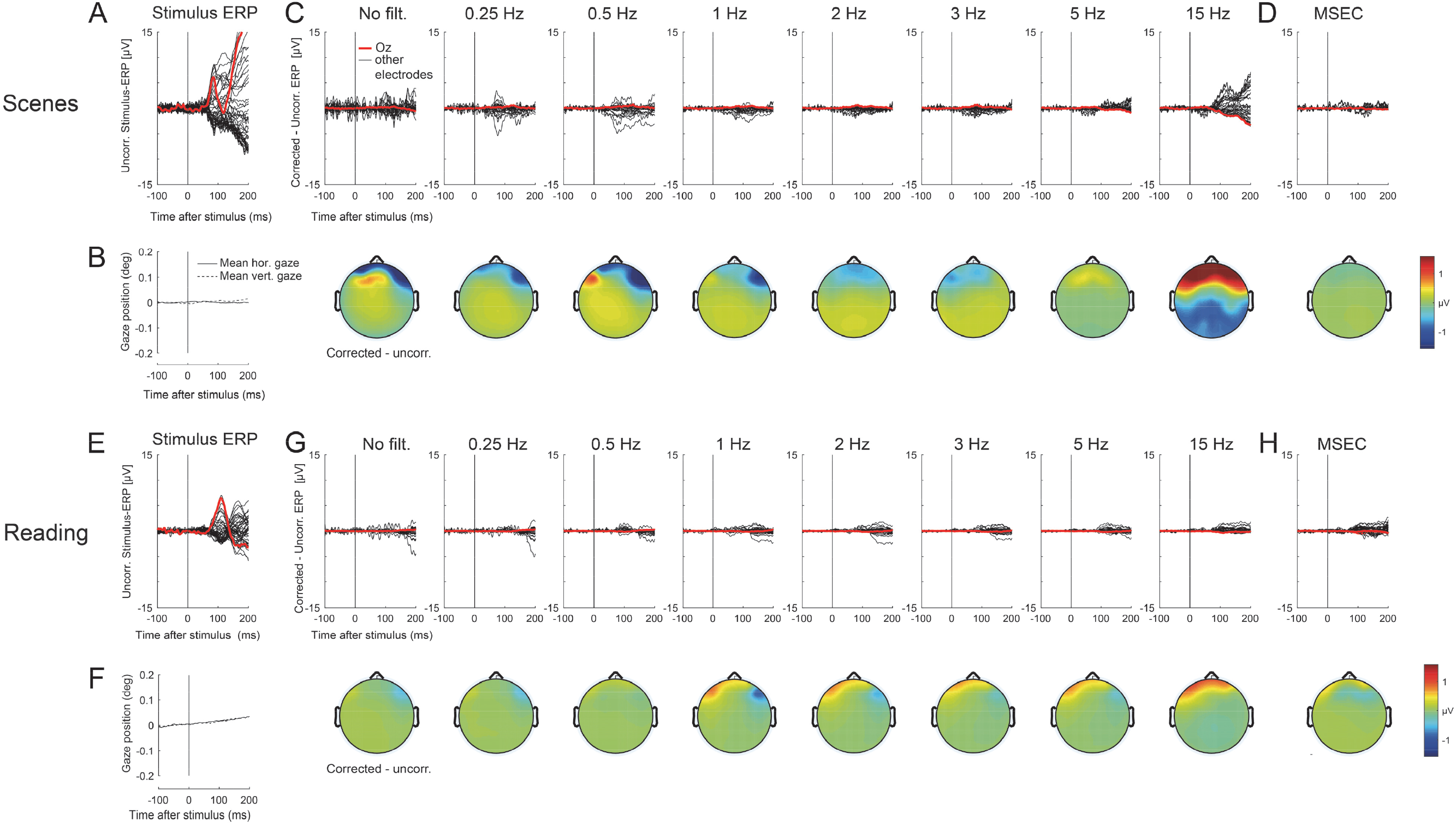
Effect of high-pass filtering on overcorrection. Shown is the stimulus-locked ERP for (virtually) eye movement-free epochs. (**A, E**) Grand-average ERPs in the *Scene* (**A**) and *Reading* (**E**) experiment. Time zero marks image/sentence onset, respectively. (B, F) Averaged horizontal and vertical eye-track. (**C, G**) Difference waves of the stimulus-ERP after ocular correction minus that without any correction for ICAs trained on differently filtered data. Scalp maps depict the corresponding difference topography between 0-200 ms. In many cases, ICA introduced overcorrection at frontal electrodes. (Note: ICAs shown here were computed on training data with overweighted SPs, low-pass filtered at 100 Hz). (**D, H**) Corresponding results for MSEC.

A first interesting finding is that all ICA variants produced some changes in eye movement-free intervals, regardless of filter settings. Crucially, however, overcorrection depended again strongly on details of the preprocessing, as confirmed by a main effect of *high-pass filter, F*(19,418) = 13.214, ɛ = 0.098, *p <* 0.0001, ηG^2^ = 0.208. More precisely, overcorrection and high-pass filtering displayed a U-shaped relationship such that with increasing cutoffs, distortions of the stimulus-ERP first decreased, then reached a plateau (from about 2 to 5 Hz), and then rebounded again. This effect of *high-pass filtering* also interacted with that of *overweighting*, *F*(19,418) = 6.43, ɛ = 0.280, *p* < 0.0001, ηG^2^ = 0.014.

Maps in Figure 7 show the scalp distribution of overcorrection for some ICA solutions. Distortions occurred mostly at frontal channels and their topographies closely resembled those of typical CR and/or SP artifacts in most cases, suggesting that the activity time courses of some of the removed ocular ICs were non-zero during eye movement-free intervals.

### Comparison to MSEC

Figures 3, 4, 5, and 7 also contain the corresponding results for MSEC. To assess its performance, MSEC was compared to a near-optimal ICA variant (2-100 Hz, with overweighting) using paired t-tests. In terms of CR artifact removal, MSEC performed not significantly better or worse than this optimized ICA (*p* = 0.86 for *Scenes*, *p* = 0.13 for *Reading*). However, MSEC produced less overcorrection than the optimized ICA variant for *Scenes*, *t*(11)=2.41, *p* < 0.05, although the actual numerical difference in overcorrection was small (Figure 3C). For Reading, overcorrection did not differ between methods. Importantly, MSEC failed to remove the SP (Figure 5), which was only reduced to a third of its original amplitude, and remained significantly larger than with optimized ICA, both for *Scenes*, *t*(11) = 8.65, *p* < 0.0001 and *Reading*, *t*(11) = 9.6, *p* < 0.0001.

### Choice of threshold

All analyses above were based on a threshold of 1.1. At this threshold, the criterion removed on average of 5.3 components for *Scenes* (SD = 1.37) and 4.0 components for *Reading* (SD = 0.74), a difference that is likely explained by the absence of vertical saccades in reading. Figure 8 summarizes effects of changing the threshold. For this analysis, the other parameters wer e fixed at near-optimal values (bandwidth 2-100 Hz, with overweighting). Ideally, we would want a bimodal distribution of variance ratios that cleanly separates between ocular and non-ocular ICs. As Figure 8A shows, the observed distribution was indeed fairly bimodal, with relatively few ICs located in a “gray area” between 1.0 and 1.4. As expected, decreasing the threshold increased the number of rejected ICs (Figure 8B). Overweighting had no effect on the number of removed ICs at any threshold (Figure 8C).

**Figure 8.**
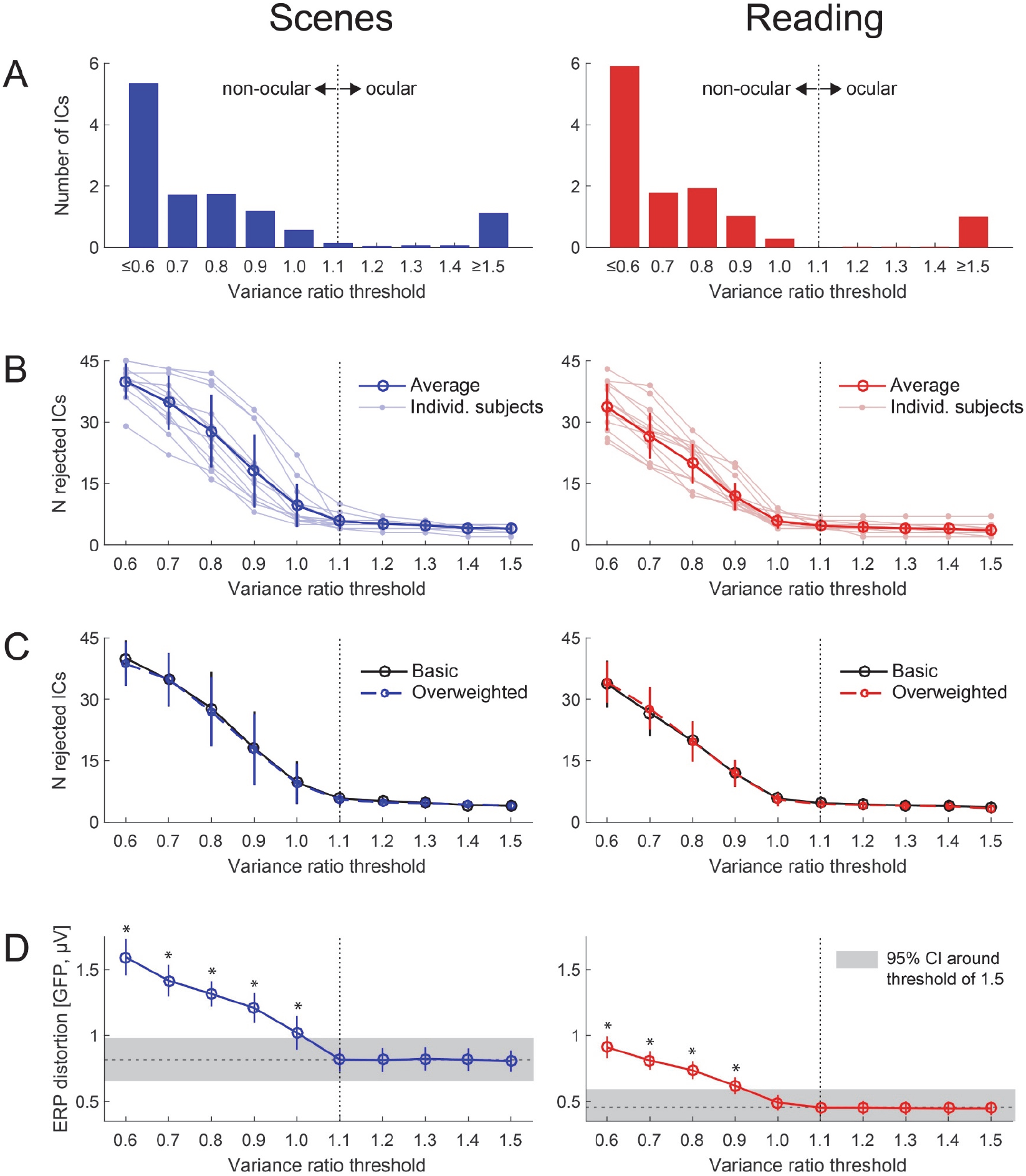
Selecting a threshold for component classification (Plöchl et al., 2012). (A) Distribution of variance ratios for all ICs, shown here for a near-optimal ICA variant (2-100 Hz, with overweighting). The threshold of 1.1, used in all analysis above is highlighted by the vertical line. For both *Scenes* (left) and *Reading* (right), there was a rather clear separation between ICs with a low vs. high variance ratio. (B) Number of ICs rejected as “ocular” as a function of threshold. (C) Overweighting SPs in the training data did not influence the number of rejected ICs. (D) Effect of threshold on overcorrection, with y-axis indicating the distortion of artifact-free epochs. Gray shading marks the 95% confidence interval (CI) around the threshold of 1.5. Compared to this lenient threshold, overcorrection increased significantly (*p* < 0.05, asterisks) once the threshold was lowered beyond 1.1 (for *Scenes*) or 1.0 (for *Reading*), indicating that 1.1 is a suitable threshold to avoid overcorrection (while ensuring good CR and SP correction).

Figure 8D shows the relationship between threshold and overcorrection. Whereas overcorrection was not significantly different for thresholds between 1.5 and 1.1, it increased once the threshold was lowered further. Specifically, for *Scenes*, overcorrection started to increase (relative to a lenient threshold of 1.5) once the threshold was lowered to 1.0 or less (comparison threshold 1.0 vs. 1.5; *t*(11) = 4.97, *p* < 0.001). For *Reading*, a statistically significant increase happened at thresholds of 0.9 or less, *t*(11) = 2.90, *p* < 0.05; however, as the right panel of Figure 8D shows, a numerical increase was already seen at 1.0. Together, these results indicate that 1.1 is indeed the lowest suitable threshold that avoids overcorrection, while also almost fully removing ocular artifacts (given optimized training data, see above).

## Discussion

Human behavior is characterized by frequent saccades of varying amplitude and direction. Although ICA has been previously applied to reduce the strong artifacts generated during natural viewing situations (such as visual search, reading, face perception, or whole-body motion), little is known about how to adapt the procedures for this purpose. Here I used eye-tracking to explore the parameter space for data preprocessing, to select ocular component, and to objectively validate correction outcomes. Specifically, the goal was to combine the concepts of optimized filtering (Winkler et al., 2015), artifact overweighting (e.g. Keren et al., 2010), eye tracker-guided component classification (Plöchl et al., 2012), and eye tracker-based quality control (Dimigen et al., 2011) into one pipeline. Results show that if parameters are optimized, Infomax ICA can remove virtually all artifacts from natural scene viewing and reading with little overcorrection. Below, I first discuss the individual parameters and then summarize specific recommendations.

### High-pass filter

Adverse effects of slow signals on unmixing quality are known among ICA practitioners (see also Miyakoshi, 2018) and many filter their input data at cutoffs between 0.1 and 1 Hz (e.g. Nieuwland et al., 2018). Similar settings have also been used in SRP/FRP research (e.g. 0.5 Hz; as used in Henderson et al., 2013; Nikolaev et al., 2011; and Ossandon et al., 2010). However, filter effects have rarely been formally investigated (Winkler et al., 2015) and not in tasks with multiple eye movements, where filtering was hypothesized to be crucial. Results confirm this hypothesis. Whereas the impact of inadequate filtering on the resulting ERP waveforms was comparatively mild in the oddball task investigated by Winkler and colleagues (cf. their Figure 5), effects were strong for *Scenes* and *Reading*. Without any filtering (i.e. 10 s time constant), ICA produced the strongest distortion of eye movement-free intervals and also left large residual CR artifacts in the saccade-related potentials; however, even at the commonly used cutoff of 1 Hz, correction was still suboptimal. Instead, filtering at 2 or 2.5 Hz produced the smallest residual CR artifacts and the least overcorrection for scene viewing. For sentence reading, the numerically best cutoff was even higher (at ~4 Hz), but filtering at 2.5 Hz still produced good results. Although the interaction of *High-pass filter* with *Experiment* did not reach significance (after sphericity correction), this pattern is likely explained by the stronger summation of CR artifacts from repeated rightward saccades during reading and indicates that aggressive filtering is especially important in paradigms with an assymetric distribution of saccade orientations. If the cutoff was raised much higher, beyond about 7.5 Hz, residual CR artifacts and overcorrection increased again markedly (in line with Winkler et al., 2015).

Of course, it should be noted that the most suitable cutoff frequency not only depends on filter steepness (here I used the frequency-dependent defaults in EEGLAB), but likely also on the length of the epochs entered into ICA. All filtering in the present study was done in addition to mean-centering the epochs, which were 3 s long (since this would be long enough for a time-frequency analysis; Miyakoshi, 2018). Since mean-centering also suppresses low frequencies (Groppe et al., 2009), optimal cutoffs may differ for extremely short epochs.

### Low-pass filter

Another parameter of interest was the upper cutoff frequency which is often set to 30-45 Hz for traditional ERP analyses. Here I found that SP correction was significantly better if filters were kept open to 100 Hz (Figure 5), even though this meant that the input data contained more line noise and scalp EMG. The likely reason for this benefit is that the spectrum of the SP extends well beyond 45 Hz (see also Figure 6), meaning that this artifact is larger – and presumably better modeled – without low-pass filtering. Of course, without low-pass filtering, a large proportion of the produced ICs may reflect scalp EMG. This is not a problem if the goal is to remove ocular ICs and then back-project the data. If the experimenter plans to work with the neural ICs in source space instead (and wants to avoid problems associated with re-running ICA on rank-reduced data after removing EMG sources; Artoni, Delorme, & Makeig, 2018), it may be better to keep the low-pass filter and only optimize the other two parameters (high-pass and overweighting), since this may yield more neural components (at the cost of a slightly less clearly modeled SP).

### Overweighting spike potentials

In combination with suitable filtering, spike potential overweighing massively improved SP correction and allowed to remove this artifact – and its associated distortions in the beta and gamma band – from both paradigms (Figures 5 and 6). These findings are consistent with the beneficial effects of overweighting for suppressing gamma-band artifacts from much smaller microsaccades in steady-fixation experiments (Keren et al., 2010; Craddock et al., 2016). Overweighting did not influence the number of removed ICs, suggesting that it produced qualitatively better SP components. I also observed that overweighting was only effective if the peri-saccadic epochs taken from the basic training data were again mean-centered across their brief 30 ms duration before appending them, most likely because this further emphasizes signal variance in high frequency bands. In line with this observation, results in Figure 3B show that both overweighting and low-pass filtering became irrelevant if the training data was already high-pass filtered at extreme cutoffs (e.g. 10 Hz), because in such data, the SP accounts for sufficient variance. Importantly, however, such extreme filtering produced bad CR correction and strong overcorrection. The tradeoff to remove *both* the SP and the CR artifact is therefore to combine moderate high-pass filtering (e.g. at 2-2.5 Hz for scenes) with the overweighting of SPs and no low-pass filtering.

In the current study, the appended samples constituted 50% of the training data. Since overweighting increases the length of the training data and therefore ICA computation time, it raises the question of how many samples need to be added. *Appendix B* presents a supplementary analysis in which the proportion of SP samples varied between 5% and 50% of the total training data. In this analysis, SP correction improved with increasing overweighing proportions at least up to 50%, in particular if the high-pass filter was chosen poorly (e.g. 0.1 Hz).

### Overcorrection & threshold choice

One question motivating the current study was whether ICA distorts neurogenic signals. Results show that all tested ICA variants modified the EEG during intervals free of significant oculomotor activity. In the scene viewing data, ICA also produced stronger distortions that MSEC, which was explicitly designed to reduce overcorrection (Berg & Scherg, 1994). Interestingly, distortions mostly affected frontal sites and their scalp distributions resembled those of SP and CR artifacts (Figure 7). Given this topographical resemblance, one might suspect that these changes in the stimulus-ERP were not really caused by “overcorrection”, but simply reflect the removal of tiny, overlooked artifacts elicited by undetected microsaccades (Meyberg et al., 2017; Plöchl et al., 2012), drift, microtremor (Onton & Makeig, 2009), or even miniature eye lid movements (see Footnote 1). While it is true that the eyes are never motionless (Rolfs, 2009), this seems unlikely for at least two reasons: First, it is unlikely that the sensitive detection algorithm (Engbert & Kliegl, 2003) missed a relevant number of microsaccades; and significant binocular drift was also eliminated. Second, and more importantly, if this was the case, overcorrection should have been largest at filter settings that were also the most effective at removing artifacts. However, the opposite was true: “Bad” ICA solutions – those that failed to suppress all CR and SP artifacts – generated the *strongest* changes in eye movement-free intervals, whereas “good” ICA solutions – those that effectively removed ocular artifacts (see Figure 3A and 3B) – produced the *least* overcorrection (Figure 3C). The more likely explanation is therefore that with a suboptimal unmixing, some neural activity or noise was modeled in the activity time courses of the ocular ICs that were later removed from the data, thereby also affecting eye movement-free intervals.^3^

As expected, the distortion of stimulus-ERPs increased if the variance ratio threshold was set too low; that is, once it was lowered to 1.0 (for *Scenes*) or 0.9 (for *Reading*). Thus, a threshold of 1.1, as initially proposed by Plöchl et al. (2012), appears to be a suitable choice.

### Comparison to MSEC

Surrogate MSEC (Berg & Scherg, 1994) was used as a benchmark for Infomax ICA. In terms of CR correction and overcorrection, this alternative method performed as well or better than the best ICA solutions obtained here. Crucially, however, at least with its default procedures (Scherg, 2013; see *Appendix A*), MSEC failed to remove the SP. MSEC also has practical drawbacks: First, it requires the experimenter to record 5-10 min of isolated eye movements from each participant before or after the experiment. Second, to my knowledge, the method is currently only implemented in proprietary software (see Berg, 2003 for an outdated open software). Finally, it provides a less flexible framework than ICA for removing non-ocular artifacts. Nevertheless, MSEC appears to be a viable alternative if the focus is on removing CR artifacts with little brain-signal distortion.

### Other challenges when analyzing multi-saccadic EEG

Artifacts are only one of four challenges when analyzing experiments with multiple saccades (Dimigen et al., 2011). The other problems relate to the (1) integration of eye-track and EEG, (2) the temporally varying overlap between the neural responses generated by successive fixations, and (3) the complex influences of visual and oculomotor low-level variables (such as saccade size) on the morphology of the post-saccadic lambda waves. The first of these problem can now be solved with dedicated toolboxes (such as EYE-EEG, see also Baekgaard, Petersen, & Larsen, 2014; Xue, Quan, Li, Yue, & Zhang, 2017), whereas the latter two are effectively addressed by analyzing the artifact-corrected EEG with regression-based linear deconvolution models (Burns, Bigdely-Shamlo, Smith, Kreutz-Delgado, & Makeig, 2013; Dandekar, Privitera, Carney, & Klein, 2011; Ehinger & Dimigen, 2018; Kristensen, Rivet, & Guérin-Dugué, 2017; Smith & Kutas, 2015). If ocular correction is also optimized, there are now viable solutions to all four problems.

### Conclusions & recommendations

Results motivate the following recommendations: First, ICAs of free viewing data should be trained on data high-pass filtered around 2-2.5 Hz (possibly higher for reading). Second, frequencies > 40 Hz should remain in the data, since this improves SP correction. Third, correction is further improved if SPs are overweighted. Fourth, a threshold of 1.1 is suitable for component identification (in combination with a saccade window that begins −10 ms before saccade onset), with lower thresholds leading to overcorrection. Infomax ICA trained in this manner removed almost all artifacts with comparatively little distortion of neural activity and no need for any subjective classifications by the experimenter. Since the suitable parameters were overall similar for scene viewing and reading, they probably also generalize to other free viewing taks, e.g. in virtual reality or during mobile brain/body imaging (Gwin et al., 2010).

Finally, it should be noted that although the parameters explored here are likely among the more important ones, they only cover some of the choices when designing the ICA pipeline. Parameters not investigated include the specific algorithm used (e.g. Infomax vs. AMICA), the overall number of data points (*k*) submitted to ICA, or the question whether and how the variance ratio criterion should be combined with other flagging methods (Chaumon et al., 2015). The eye tracker-based quality measures proposed here may help to quantify the role of these other parameters in the future.

### Implementation

A function to create training data with overweighted artifacts was added to the EYE-EEG extension for EEGLAB and a simple Matlab script implementing the current procedures is found at www.github.com/olafdimigen/opticat.

## Appendix

## Appendix A Surrogate Multiple Source Eye Correction

Surrogate MSEC separates ocular artifacts from neural activity based on spatial definitions provided by two sets of scalp topographies: artifact topographies and brain-signal topographies. A set of artifact topographies is defined empirically by averaging stereotypical eye movements recorded from each participant during a short “calibration” session immediately before or after the actual experiment. During this 5-10 min session, participants executed 30 self-paced saccades with an amplitude of 15° in each of the four cardinal directions (left, right, up, down) plus about 30 spontaneous eye blinks during central fixation. Following the BESA handbook (Scherg, 2003), the EEG data of this calibration session was then band-pass filtered from 0.5-8 Hz and epochs were cut around eye movement onsets. The average across the 400 ms interval following eye movement onset was used as the artifact topography for this type of movement. To remove saccade-related neural activity from these “prototypical” individual artifact topographies, epochs were then entered into a PCA in BESA. For the current study, only the first spatial primary component (factor), which typically explained 90-99% variance, was retained to define the scalp topography for each of three types of artifact (horizontal saccade, vertical saccade, and blink).

In addition, a set of “typical” brain topographies was defined by the scalp projections of an equivalent dipole source model in BESA. This generic (“surrogate”) brain model was the same for all participants and defined by BESA standard model “*BR_Brain Regions_LR.bsa*”, which contains 15 regional sources (each consisting of three orthogonal dipoles) that are widely distributed across the head. The regularization constant for brain activity was set to the default of 2%.

To perform the actual correction, activity time courses (i.e. “source waveforms”) for the artifact topographies are estimated by inverting a matrix (see Ille et al., 2002, p. 123) that contains as columns the three empirically defined artifact topographies (horizontal, vertical, and blink) plus the brain topographies defined by the generic source model. In a final step, the source waveforms estimated for the three artifact topographies are subtracted from the raw EEG, yielding a corrected version of the data. Note therefore, that with surrogate MSEC, brain activity in the experiment is *not* directly modeled; instead the “typical” brain topographies defined by the dipole model are only added to the matrix to reduce the subtraction of neural activity that is spatially correlated to the artifact topographies. In other words, the purpose of the dipole model is to reduce overcorrection. For the current study, I followed procedures in the BESA handbook (Scherg, 2013), which are also largely identical to those used in previous free viewing studies (Dimigen et al., 2011, 2012).

Finally, please note, that it might be possible to improve SP correction if the recommended MSEC procedures (see Scherg, 2013) were modified such that (1) the EEG data from the calibration session is not low-pass filtered at 8 Hz, (2) the post-saccadic time window on which artifact topographies are computed is shrunk and shifted backwards so that it focuses more on the peri-saccadic interval and (3) several spatial PCA factors, rather than just the first one, are retained as artifact topographies for each type of eye movement (because the SP may be captured in one of the PCA factors explaining less variance).

## Appendix B Amount of overweighting

**Figure A1.**
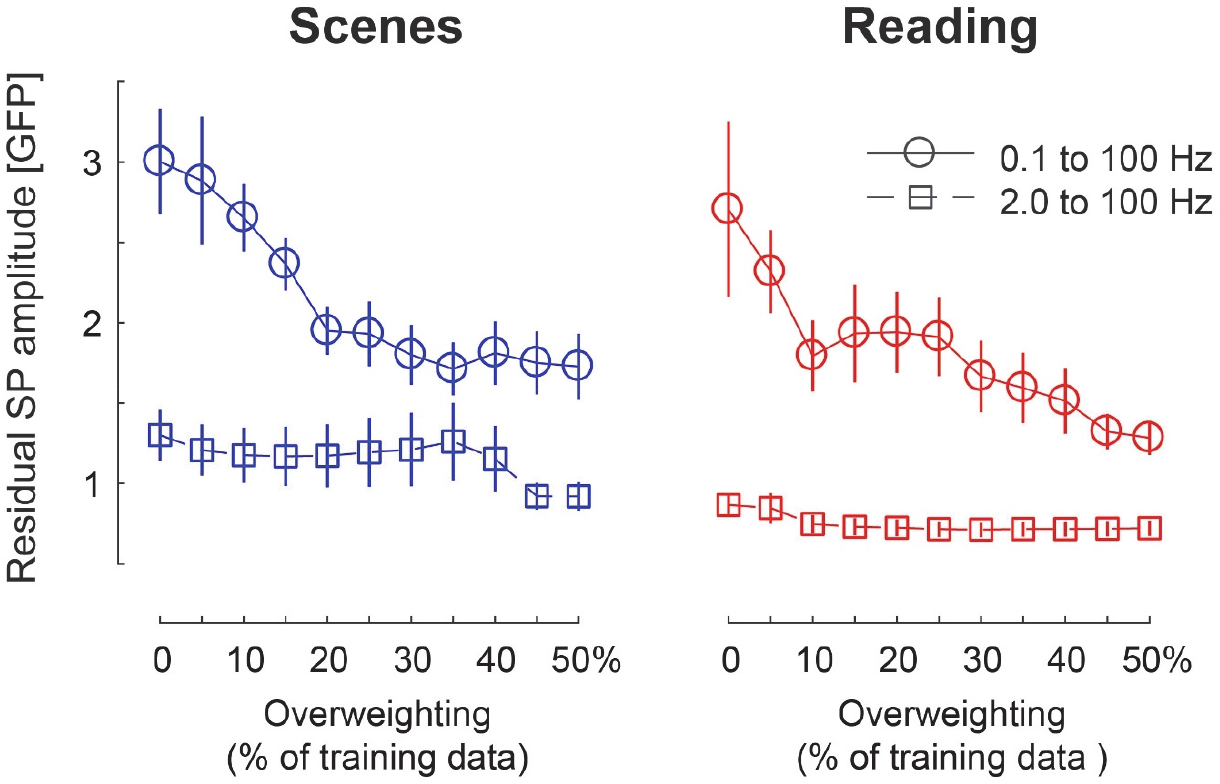
Relationship between the proportion of peri-saccadic samples in the total ICA training data and the size of the SP artifact after correction. The relationship is shown here for training data filtered from 0.1 to 100 Hz (round markers) and 2 to 100 Hz (square markers). 0% means no overweighting whereas 50% means that the length of the training data was doubled by appending SP samples. Correction was improved by adding more SPs, especially if the high-pass filter was chosen poorly (here: 0.1 Hz).

## Appendix C Correction of vertical saccades (scene viewing only)

**Figure A2.**
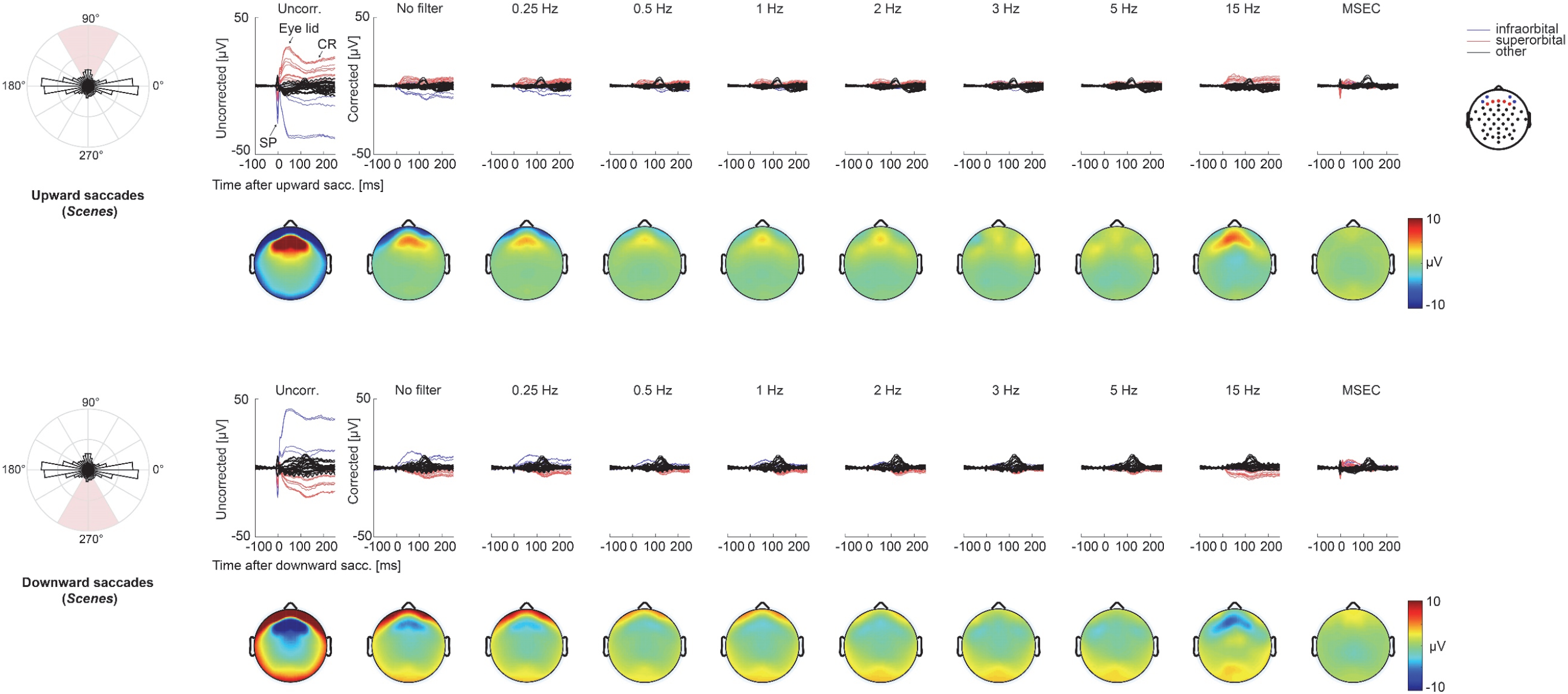
Effect of high-pass filtering on correction of upward (top panels) and downward (bottom panels) saccades (±30°) in the *Scenes* experiment. For a general description of this plot refer to Figure 4. Blue lines in this plot indicate four peri- and infraorbital EOG electrodes. Red lines mark six prefrontal channels (Fp1, Fpz, Fp2, AF7, AFz, AF8). Results were overall similar to those obtained for the much more frequent horizontal saccades. However, as this figure shows, the most suitable low-pass filter setting was slightly higher for vertical (3 Hz) than horizontal (2 to 2.5 Hz) saccades. Results shown in this plot are for ICAs trained on data filtered at 100 Hz with overweighted SPs.

## Footnotes

1 Blinks can be objectively identified with eye-tracking because pupil and corneal reflex tracking is lost during blinks. An unlikely possibility that cannot be fully excluded is the existence of smaller twitches of the eye lids during cognitive task, which may not occlude the pupil, but still create small blink artifacts. Furthermore, even during microsaccade-free intervals, the eyes are never entirely motionless but show slow (< 0.5°/s, Rolfs, 2009) conjugate or non-conjugate *drift* movements of limited spatial extent (i.e. resembling a random walk) as well as a binocularly synchronized, high-frequency (~30-100 Hz, Rolfs, 2009) *microtremor* of extremely small (~0.1 to 0.5 min-arc) amplitude. Significant conjugate drift was eliminated by the criteria applied here. Regarding microtremor, it has been putatively suggested by Onton & Makeig (2009) that it produces a tiny, high-frequency (> 40 Hz) oscillation in peri-ocular EEG channels; however, such a small continuous oscillation would not have affected the overcorrection measure used here in a significant manner.

2 The SP begins before saccade onset. A comparison of 20 different saccade window sizes (not reported here) showed that this definition (−10 ms before saccade until saccade offset) distinguished best between ocular and non-ocular ICs (i.e. maximized the distance between the two variance ratio distributions separated by a threshold of 1.1; see also Figure 8A).

3 The fact that the distortion of artifact-free intervals was not larger for filtered (e.g. at 2 Hz) than unfiltered training data also indicates that it is permissible to transfer ICA weights trained on filtered data to the unfiltered version of the same data (e.g. Viola et al., 2010).

